# Drug-free nasal spray as a barrier against SARS-CoV-2 infection: safety and efficacy in human nasal airway epithelia

**DOI:** 10.1101/2021.07.12.452021

**Authors:** Fabio Fais, Reda Juskeviciene, Veronica Francardo, Stéphanie Mateos, Samuel Constant, Massimo Borelli, Ilja P. Hohenfeld, Thomas Meyer

**Affiliations:** Altamira Medica AG, Zug, Switzerland; Texcell SA, Evry, France; Epithelix Sarl, Geneva, Switzerland; Life Sciences and Technologies Department, School of PhD Programmes, Magna Graecia University, Catanzaro, Italy

## Abstract

**Background:** For SARS-CoV-2 and other respiratory viruses, the nasal epithelium is a key portal for infection. Therefore, the nose is an important target of prophylactic and therapeutic interventions against these viruses. We developed a nasal spray (AM-301, a medical device marketed as Bentrio) to protect against infection by SARS-CoV-2 and potentially other viruses.

**Aims of the study:** To test the safety and efficacy of AM-301 against SARS-CoV-2 infection.

**Methods:** AM-301 was tested on an in vitro 3D model of primary human nasal airway epithelium. Safety was assessed in assays for tight junction integrity, cytotoxicity and cilia beating frequency. Efficacy against SARS-CoV-2 infection was evaluated in prophylaxis and infection mitigation assays.

**Results:** AM-301 did not have any detrimental effect on the nasal epithelium. Prophylactic treatment with AM-301 reduced viral titer significantly vs. controls over 4 days, reaching a maximum reduction of 99%. When treatment with AM-301 was started 24 or 30 h after infection, epithelia that received the formulation had a 12- or 14-fold lower titer than controls.

**Conclusion:** AM-301 was found to be safe in vitro, and it significantly decelerated viral titer growth in experimental models of prophylaxis and mitigation. Its physical (non-pharmaceutical) mechanism of action, safety and efficacy pave the way for further investigation of its possible use against a broad spectrum of viruses, allergens and pollutants.

## Introduction

The COVID-19 pandemic has had a massive toll on life [1] and has strained the capacity of healthcare institutions [2–4]. The pandemic continues to have a major impact on daily conduct in our attempt to prevent the spread of the causative agent, severe acute respiratory syndrome coronavirus 2 (SARS-CoV-2). This virus is the third coronavirus known to cause severe disease in humans, and its emergence follows two global outbreaks by SARS in 2002-2003 and the Middle East respiratory syndrome-related coronavirus in 2012 (reviewed in [5]). To arrest the current pandemic, governments worldwide have variably implemented a series of non-pharmaceutical interventions, from social distancing, school closures and travel restrictions to improved hygiene and face mask requirements [6,7]. Despite ongoing vaccination campaigns, there is a pressing need for new, effective personal protection measures against the infection.

SARS-CoV-2 infection is mainly contracted from airborne virions [8,9]. Upon inhalation, the virions bind cells that express angiotensin-converting enzyme 2 (ACE2), and after host cell proteases cleave the viral spike protein, the virions enter cells and cause infection [10]. Therefore, the main site of initial infection is the nasopharyngeal epithelium and, in particular, the ciliated cells, which express high levels of ACE2 and the proteases TMPRSS2 and furin on their apical side [11]. During the first days of SARS-CoV-2 infection, the nasal tract has a high viral load [11,12]. It is during this period that an infected person is most contagious towards others [13] and at risk of developing, by aspiration along the naso-oropharynx-lung axis [14], a lower respiratory infection that may proceed to pneumonia and severe COVID-19. The nasal microenvironment, therefore, plays a major role in SARS-CoV-2 infection and clinical course (reviewed in [15]).

The airway epithelium of the nasal mucosa protects from infection by mainly serving as a physical barrier through the production of mucus, which traps pathogens, and the clearing action of cilia, which discharge the mucus into the nasopharynx from where it is eventually swallowed [16]. A second line of protection is provided by immune cells resident in the nasopharynx-associated lymphoid tissue [15]. Together, mucociliary clearance and immune responses should protect the nasal epithelium from pathogens, but infection can ensue in cases of high viral exposure or dysfunction of these mucosal defenses. Therefore, there is interest in developing simple and safe interventions to enhance nasal barrier functioning.

Several nasal sprays are commercially available for the treatment of respiratory infections such as the common cold and influenza, and recent research has investigated the possibility of repurposing these sprays as COVID-19 prophylactic agents. One study [17] found that two products containing carrageenan, but no pharmaceutically active ingredients, inhibited SARS-CoV-2 infection in an in vitro model of the human airway epithelium. This same study found that sprays with other inert or active ingredients were cytotoxic. Carrageenans are polyanionic, sulfated polysaccharides from red seaweed; they are widely used in pharmaceutical formulations and have known virus-binding properties [18]. Other natural substances with broad pharmaceutical applications and virus-capturing properties are clays, including bentonite [19–22]. Bentonite is a clay mineral composed of thin aluminum silicate sheets with a net negative charge; these properties contribute to its ability to adsorb viral particles and molecules such as drugs [21]. We therefore hypothesized that a bentonite-containing nasal spray could protect against SARS-CoV-2 and other airborne pathogens. Bentonite suspensions can have thixotropic properties, that is, they reversibly change from a gel when undisturbed to a fluid colloid when agitated [23]. We envisioned a bentonite-containing nasal spray that could be applied as a liquid, but in the nasal cavity it would form a durable, protective gel barrier.

We therefore devised a nasal spray (AM-301, Bentrio) with the aim of providing a safe and effective means of self-protection against exposure to harmful airborne particles. Seeking to minimize potential side effects and facilitate frequent and compliant use, we selected only inert ingredients such as pharmaceutical excipients and substances that are generally recognized as safe. Because the components are neither metabolized nor absorbed and the mechanism of action is physical, not chemical, AM-301 is a non-pharmaceutical medical device. This study tested the ability of AM-301 to prevent SARS-CoV-2 infection or mitigate existing infection in vitro using a model of human nasal airway epithelium.

## Materials and methods

### Virus and cell cultures

The SARS-CoV-2 strain 2019-nCOV/Italy-INMI1 was obtained from the National Institute for Infectious Diseases Lazzaro Spallanzani IRCCS (Rome, Italy) [24]. The strain was propagated on VERO cells, collected, aliquoted, and stored at -70 °C until use at Texcell (Evry, France). For viral titration assays, VERO cells were cultured in DMEM supplemented with 4% fetal bovine serum (FBS). The cell line had been obtained by Texcell from the Pasteur Institute (Paris, France). Testing for mycoplasma contamination using the MycoTOOL Mycoplasma Real-Time PCR Kit (Roche) was negative.

MucilAir Pool tissue cultures were obtained from Epithelix (Geneva, Switzerland) [25–27]. MucilAir Pool consists of human airway epithelial cells collected from 14 healthy donors (male and female) and reconstituted as a 3D tissue in a two-chamber system; for this study, only nasal epithelial cells were used. The cells were cultured at 37°C (humidified 5% CO_2_ atmosphere) on Costar Transwell porous inserts (0.33 cm^2^ each, 24-well plates) with MucilAir serum-free culture medium (cat. no. EP04MM, Epithelix). Approximately one month after seeding, when the cultures were fully differentiated and pseudo-stratified, with basal cells, ciliated cells and mucus cells, they were exposed to air on the apical surface and were considered suitable for experimental use. Prior to testing, MucilAir Pool cultures (hereafter “MucilAir inserts”) were cultured with 0.7 mL culture medium in the basolateral chamber and air at the apical surface; the medium was changed every 3 days. Each insert had approximately 500,000 cells.

### Nasal spray formulation

The nasal spray tested in this study is a medical device containing bentonite (magnesium aluminum silicate) in a matrix composed of mono-, di- and triglycerides, propylene glycol, xanthan gum, mannitol, disodium EDTA, citric acid and water. All components are listed in the Inactive Ingredient Database of the US Food and Drug Administration (FDA), are classified as “generally recognized as safe”, or are approved for use as food additives by the FDA. The formulation is a white to light beige, aqueous gel emulsion with a pH of 6.0. AM-301 is odorless and tasteless. When we applied it to our own nasal mucosa or palmar skin, it had a soothing non-irritant, lotion-like consistency.

Before testing, AM-301 and its matrix were brought to room temperature and vigorously agitated for 10 s. The amount of product tested on MucilAir inserts was calculated to roughly correspond to the amount that would be delivered to the nostril by a standard, commercial spray applicator (140 μL per actuation). Considering that the total nasal cavity surface area is 160 cm^2^ [28] and that a nasal spray coats the anterior third of the cavity [16], then 5.3 μL of product should be tested per square centimeter of tissue. Since MucilAir inserts are 0.33 cm^2^, 10 μL of a 1:5 aqueous dilution of AM-301 or its matrix (diluted water immediately before use) was tested per insert.

### Safety assays

AM-301 was tested for potential cytotoxic effects in a series of tissue culture assays used routinely for assessing the quality of MucilAir preparations. These assays, done by Epithelix, included an assay for transepithelial electric resistance (TEER), which measures tight junction integrity [29]; an assay for lactate dehydrogenase (LDH) release into the basolateral medium, a standard measure of cytotoxicity; and an assay for cilia beating frequency (CBF), which is an index of the main function of airway cells, namely mucociliary clearance [30]. The three assays were done simultaneously on one set of 12 MucilAir inserts over a 4-day protocol with repeated apical application of the product, as described below and illustrated in **Figure 1A** (see “Tissue culture assays” below for the individual methods).

**Figure 1.**
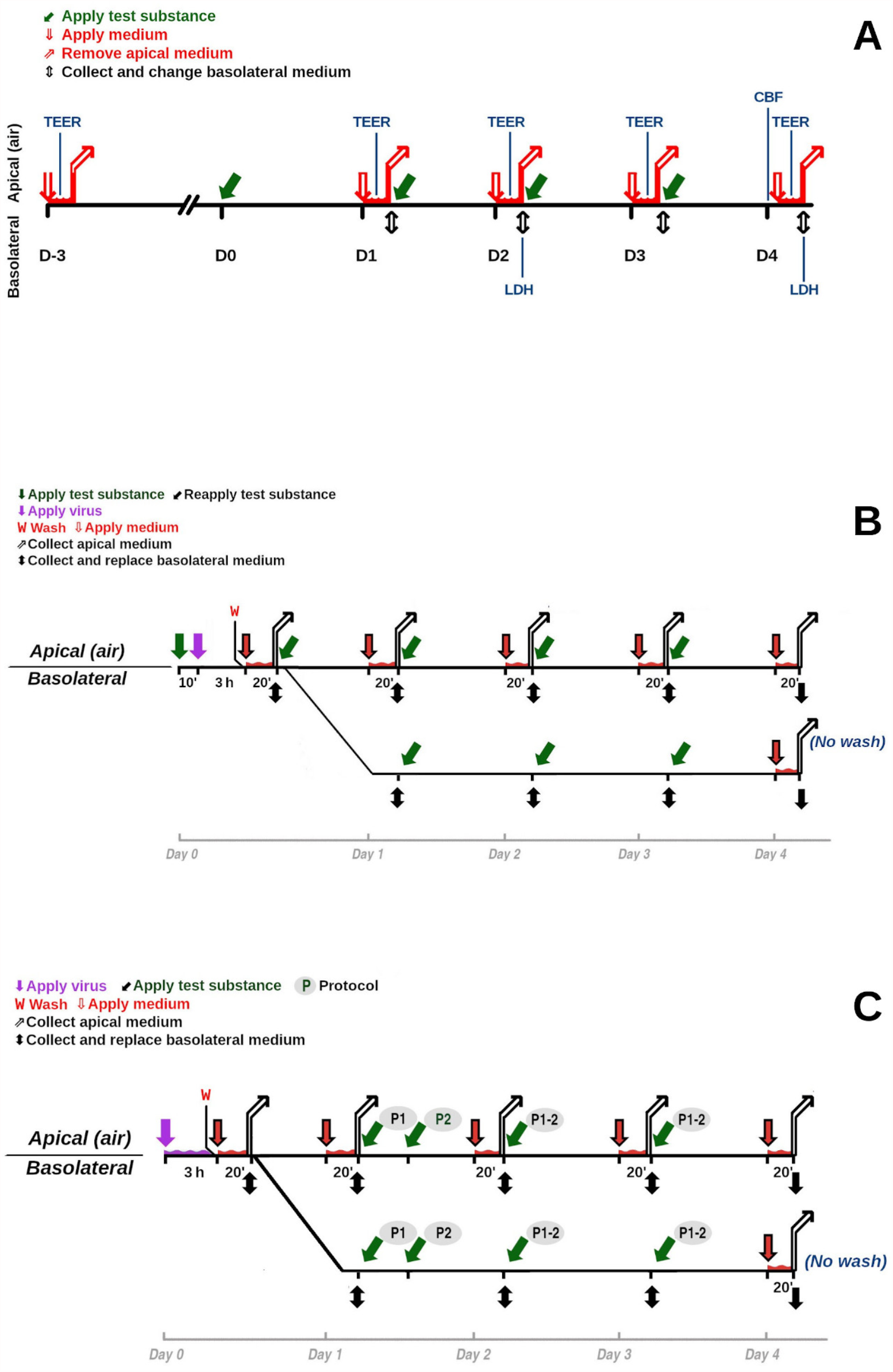
Schematics of experimental protocols. **A** Safety assays. CBF, cilia beating frequency; LDH, lactate dehydrogenase; TEER, transepithelial electric resistance. **B** Prophylaxis assay. **C** Mitigation assay.

Briefly, 3 days before use, inserts were washed apically with 200 μL culture medium (10 min), and TEER was measured to verify their integrity. The apical medium was discarded, and inserts were returned to the incubator for 3 days. On the first day of product testing (Day 0), inserts were transferred to new 24-well plate with 500 μL/well fresh medium. The apical surface was treated, in triplicate, with 10 μL of a 1:5 aqueous dilution of AM-301 or its matrix or left untreated. Inserts were returned to the incubator for 24 h.

On Day 1, 200 μL medium was applied apically for TEER measurements. The apical medium was removed, and inserts were transferred to new 24-well plate with 500 μL/well fresh medium. The inserts were treated as on Day 0 and returned to the incubator. On Day 2, 200 μL medium was applied apically for TEER measurements. The apical medium was removed, and the basolateral medium was collected for LDH assays. The inserts were transferred to a new 24-well plate with 500 μL/well fresh medium, treated as on Day 0, and returned to the incubator. On Day 3, 200 μL medium was applied apically for TEER measurements. The apical medium was removed, and inserts were transferred to new 24-well plate with 500 μL/well fresh medium; the inserts were treated as on Day 0 and returned to the incubator.

On Day 4, cilia beating on the apical surface was observed and its frequency quantified. Then, 200 μL medium was applied apically for TEER measurements. Finally, the basolateral medium was collected and assayed for LDH.

### Tissue culture assays

Transepithelial electric resistance (TEER) was measured with an EVOMX epithelial volt-ohm meter (World Precision Instruments). Briefly, 200 μL MucilAir culture medium (34°C) was applied to the apical surface of each MucilAir insert. Electrodes were washed with 70% ethanol and then with medium prior to insertion into the apical and basolateral media. Resistance values (Ω) were measured at room temperature; they were corrected and converted to TEER (Ω cm^2^) using the formula:

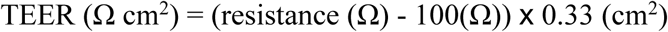

where 100 Ω is the resistance of the membrane and 0.33 cm^2^ is the surface area of the insert. The assay was performed in triplicate. Normal values for MucilAir are in the range 200–600 Ω cm^2^ [26].

Lactate dehydrogenase activity in the basolateral medium was assayed using the Cytotoxicity Detection Kit (Sigma, cat. no. 4744934001). For the high control, MucilAir inserts (n = 3) were treated apically with 100 μL lysis solution for 24 h in a tissue culture incubator and the basolateral medium was collected. For the assay, 100 μL medium from each test sample and from the high controls was transferred to a 96-well plate; culture medium alone was used as the low control (n = 3). Reaction solution was added (100 μL/well), and the plate was incubated for 15 min in the dark at room temperature. Then, stop solution was added (50 μL/well), and absorbance was read at 490 nm on a plate reader. Cytotoxicity was expressed as a normalized percentage relative to the high and low controls. Normal values for MucilAir inserts are ≤5 %, which corresponds to a physiological turnover of cells in culture [26].

Cilia beating frequency at the apical surface of MucilAir inserts was observed at room temperature under a Primovert inverted microscope (Zeiss) with a 10× objective. A Mako G-030B machine vision camera (Allied Vision Technologies) mounted on the microscope was used to capture 256 movies (125 frames per second). The registrations were analyzed with Cilia-X software (Epithelix), which calculated the cilia beating frequency. Normal values of frequency for MucilAir are in the range of 4-8 Hz.

### Efficacy assays

AM-301 was subjected to a series of in vitro efficacy assays using MucilAir inserts at Texcell. The ability to prevent SARS-CoV-2 infection was tested in a prophylaxis assay, while the ability to reduce SARS-CoV-2 replication in already infected tissues was tested in a mitigation assay. For both assays, inserts were treated apically with 100 μL of a suspension of SARS-CoV-2 in culture medium at a multiplicity of infection (MOI) of 0.5 (2.5×10^6^ TCID_50_/mL).

### Efficacy assay I: prophylaxis

In the prophylaxis assay, MucilAir inserts were apically treated with test items for 10 min before exposure to SARS-CoV-2, and viral replication was measured over 4 days with daily apical treatments (**Figure 1B**). Briefly, on Day 0, the inserts were transferred to a new 24-well plate with 500 μL/well fresh medium and washed apically by incubation with 200 μL medium for 20 min at 34°C in a humidified 5% CO_2_ atmosphere; the apical medium was removed. The apical surface was then treated with 10 μL of a 1:5 aqueous dilution of AM-301 (n = 9) or of its matrix (n = 3), or with 10 μL saline diluted 1:10 in water (n = 3), or it was left untreated (n = 3). After 10 min, without washing, the apical side was treated with 100 μL SARS-CoV-2 suspension (MOI = 0.5); viral suspension was added to all inserts except 3 treated with AM-301 and 3 untreated (negative controls). Infection was allowed to proceed for 3 h in a 34°C incubator and stopped by gentle washing of the apical surface with 200 μL medium (3 times). Viral replication was assessed immediately and daily over 4 days (n = 3 per time point) as follows:

1. *Viral sampling*. 300 μL medium was applied to the apical surface. After 20 min at 34°C, the conditioned apical medium was collected and stored at -70°C until analysis.
2. *Medium change*. Inserts were transferred to a new 24-well plate with 500 μL/well fresh medium. The conditioned basolateral medium was collected and stored at -70°C for possible future analyses.
3. *Repeat treatment*. The apical surface was treated with product or left untreated, as described above, and the inserts were returned to the 34°C incubator.

For 3 inserts treated with AM-301 (called “AM-301 no wash”), viral sampling was not done on Days 1-3. Treatments were repeated daily, and sampling was done only on Day 4.

### Efficacy assay II: mitigation

In the mitigation assay, MucilAir inserts were apically infected with SARS-CoV-2, and viral replication was measured over 4 days with no treatment (untreated, n = 3) or with daily apical treatments of physiological saline, AM-301 or its matrix (**Figure 1C**). Two experimental protocols (P1 and P2; 15 inserts each), differing in the interval between the end of viral exposure and first application of test substances, were followed.

Briefly, on Day 0, the inserts were transferred to new 24-well plates with 500 μL/well fresh medium and washed apically by incubation with 200 μL medium for 20 min at 34°C in a humidified 5% CO_2_ atmosphere; the apical medium was discarded. The apical surface was infected with 100 μL SARS-CoV-2 suspension (MOI = 0.5); viral suspension was added to 12 inserts per protocol while 3 inserts served as negative viral controls. Infection was allowed to proceed for 3 h in a 34°C incubator and stopped by gentle washing of the apical surface with 200 μL medium (3 times). Viral replication was assessed immediately by addition of 300 μL medium to the apical surface and incubation for 20 min at 34°C; the conditioned apical medium (Day 0) was collected and stored at -70°C until analysis. Inserts were transferred to new 24-well plates with 500 μL/well fresh medium and returned to the incubator.

On Day 1, viral replication was assessed as on Day 0 for all inserts per protocol except for the 3 “AM-301 no wash” inserts. All inserts were transferred to new 24-well plates with 500 μL/well fresh medium. Inserts in protocol P1 were treated with AM-301, its matrix or physiological saline, and returned to the incubator. An additional 15 inserts (protocol P2) were returned to the incubator without treatment; they were instead treated at the 30 h time point.

On Days 2 and 3, viral replication was assessed in all inserts but the “no wash” condition; the conditioned medium (Days 2 and 3) was collected and stored at -70°C. Inserts were transferred to new 24-well plates with fresh medium, treated with AM-301, its matrix or saline, or left untreated (blanks), and returned to the incubator. On Day 4, viral replication was assessed in all inserts.

### Viral titer assay

Viral titers were determined in conditioned medium collected from the apical side of MucilAir inserts. Samples were prediluted 1:32 in DMEM containing 0.5 mg/mL gentamicin (to avoid any possible interference of the test products on cell growth). Then, using one 96-well plate per sample, serial 3-fold dilutions were made in the same culture medium with 8 replicates per dilution (10 dilutions per sample); 8 wells received fresh medium (negative controls). A fixed volume (50 μL) from each well was transferred to sample titration plate, and 50 μL VERO cell suspension (10^5^ cells/mL in DMEM plus 4 % FBS) was added per well. Plates were incubated at 37°C in a 5% CO_2_ humidified atmosphere for 6 days. Then, a 0.2% crystal violet solution was added to stain DNA in live, adherent cells. The tissue culture infectious dose that killed 50 % of cells (TCID_50_) was calculated using the Spearman-Kärber method.

### Statistical analyses

Data belonging to LDH and CBF analyses were compared using the unpaired t-test; results from the TEER analyses were compared using 2-way repeated measure ANOVA followed by post hoc Tukey’s test. The level of significance was set at p<0.05. Results from the SARS-CoV-2 prophylaxis and mitigation experiments were analyzed using linear mixed-effect models with log-transformed data [31]. The minimal adequate mixed-effect model was achieved by top-down selection, adjusting for multiple comparisons. Analyses were performed using the lme4 package [32] as implemented into the programming language R [33].

## Results

### In vitro safety

MucilAir inserts were repeatedly exposed to AM-301 or its matrix (lacking bentonite), or left untreated for 4 days, and then tested in four standard assays for MucilAir integrity. TEER measurements (**Figure 2A**) at baseline were in the normal range (200–600 Ω cm^2^) for all inserts, indicating their suitability for use in toxicology testing. Mean values for inserts treated with AM-301 or matrix and for untreated inserts increased slightly over 4 days. These results indicate that the product did not have any adverse effects on tissue integrity.

**Figure 2.**
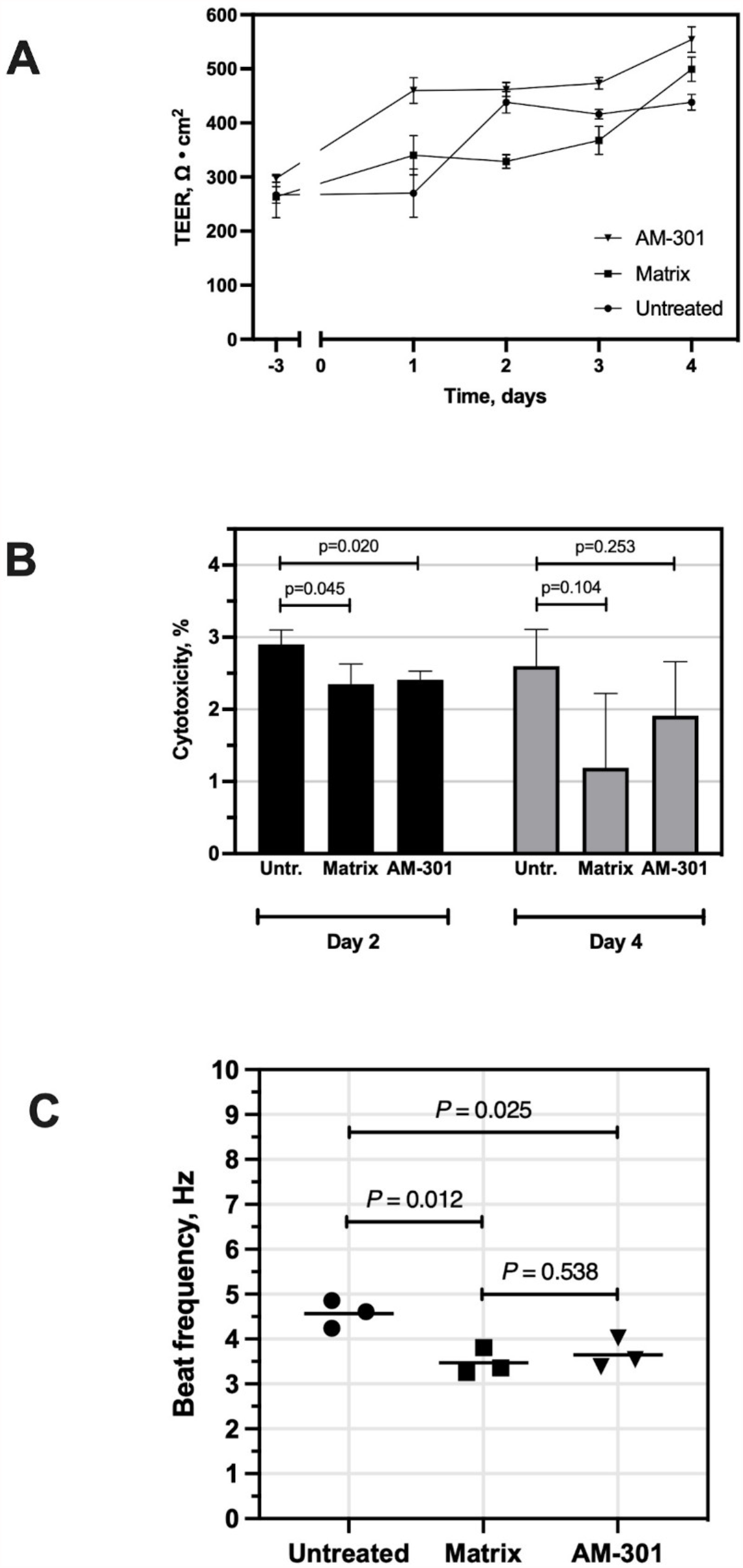
In vitro safety of AM-301 on MucilAir inserts (reconstituted human airway epithelial cells). **A** Transepithelial resistance (TEER) over 4 days of exposure to AM-301 or its matrix lacking bentonite. Significant differences were found in mean TEER by 2-way repeated measures ANOVA: Treatment, F_(2,6)_=29.30, p=0.0008; Time, F(_2.112, 12.67)_=153.0, p<0.0001; Interaction F_(8,24)_=15.76, p<0.0001. Post hoc Tukey’s test: Day –3: untreated vs. matrix, p>0.05; untreated vs. AM-301, p>0.05; matrix vs. AM-301, p>0.05; Day 1: untreated vs. matrix, p>0.05; untreated vs. AM-301, p=0.0143; matrix vs. AM-301, p=0.0264; Day 2: untreated vs. matrix, p=0.0057, untreated vs. AM-301, p>0.05, matrix vs. AM-301, p=0.0005; Day 3: untreated vs. matrix, p>0.05, untreated vs. AM-301, p=0.0049, matrix vs. AM-301,p=0.0210; Day 4, untreated vs. matrix, p=0.0476; untreated vs. AM-301, p=0.0082, matrix vs. AM-301, p>0.05. **B** Lactate dehydrogenase (LDH) release assay for cytotoxicity. Data are expressed as a percentage of the amount of LDH released by lysed cells. No significant difference between matrix and AM-301 on Day 2 (p=0.776) or Day 4 (p=0.392). **C** Cilia beating frequency after 4 days of exposure to AM-301 or its matrix.

The release of LDH to the basolateral medium, a sign of cell lysis, was assayed after 2 and 4 days of exposure to the products (**Figure 2B**). At both time points, the normalized percentage of cytotoxicity was well below 5 % for treated and untreated inserts alike. These results indicate that the product had no acute cytotoxic effects on cells, and that, in all treatment conditions, only normal cell turnover was occurring (values ≤5 %). After 4 days, cilia beating frequency at the apical surface was measured (**Figure 2C**). Untreated inserts had a mean frequency of 4.6 Hz, while inserts treated with the matrix or AM-301 had slightly lower values (3.5 and 3.7 Hz; p=0.012 and p=0.025, respectively).

### In vitro efficacy

To test AM-301’s ability to protect against SARS-CoV-2 infection of the nasal epithelium, MucilAir inserts were treated apically with the product, its matrix, or physiological saline shortly before exposure to the virus, and then treated daily for 4 days (**Figure 3A**). Viral replication was robust in both saline- and matrix-treated inserts, with more than 10-fold daily increases in titer from Day 1 to 3 and a smaller increase on Day 4. Instead, in AM-301-treated inserts, viral replication was strongly dampened. On Day 4, a 2 log reduction in TCID_50_ compared to saline control was observed, which corresponds to 99% lower viral titer. The protective effect of AM-301 was also seen in the “no wash” experiment, in which virus was allowed to accumulate over 4 days under repeated application of the product: the viral titer on Day 4 was about 13-fold lower than that observed at the same time point in both saline- and matrix-treated inserts. Viral titer for the inserts not exposed to SARS-CoV-2 was <231, indicating that the test substances and culture medium were free of viral contamination. Because viral replication was unhindered in matrix-treated samples, we can infer that bentonite in AM-301 was primarily responsible for the effect.

**Figure 3.**
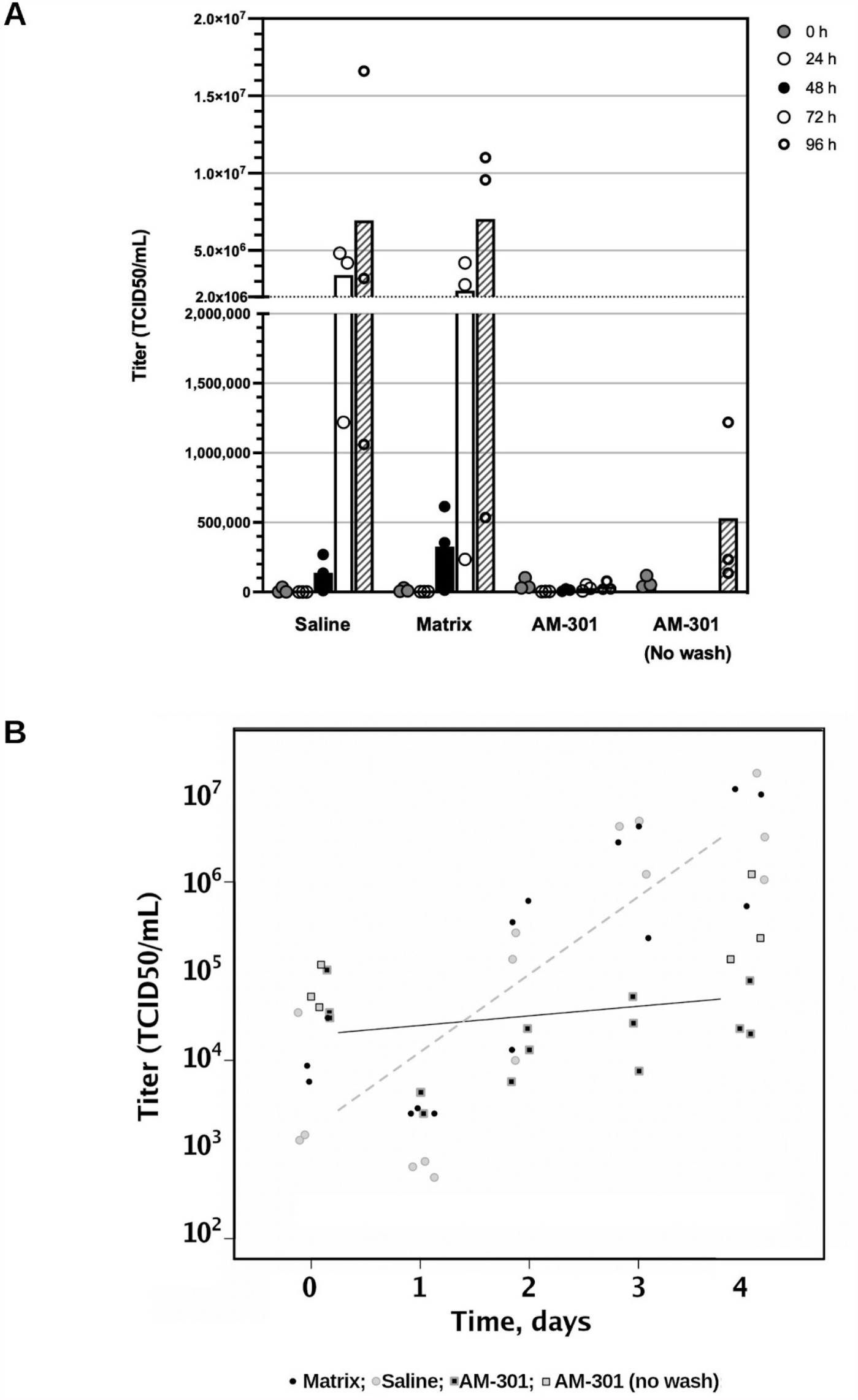
AM-301 as prophylaxis against SARS-CoV-2 infection. MucilAir inserts were treated for 10 min with physiological saline, AM-301 or its matrix, followed by viral suspension for 3 h. Then, inserts were washed and incubated for up to 4 days with daily reapplication of test substances. AM-301 (no wash): treatments were repeated daily, but viral sampling was done only on Day 4. **A** Bar chart with mean values and individual data points. **B** Linear mixed-effect model. The log-linear scatter plot shows individual log-transformed data and regression lines for negative control samples (saline- and matrix-treated inserts; dashed line) and for AM-301-treated inserts (continuous line) (t=5.49, p<0.001).

Data from the above prophylaxis experiment were statistically analyzed using a linear mixed-effect model (**Figure 3B**). The time profile of SARS-CoV-2 infection in inserts that received a prophylactic treatment with AM-301, with or without washing procedure (AM-301 and AM-301 no wash, respectively), was significantly decelerated compared to that of inserts that received saline or matrix (t=5.49; p<0.001).

To test AM-301’s ability to mitigate an existing SARS-CoV-2 infection of the nasal epithelium, MucilAir inserts were infected and then treated with test substances starting only 24 or 30 h after infection (**Figure 4**). In Protocol 1 (test substances applied 24 h after infection; Figure 4A), there was robust viral titer growth in the saline- and matrix-treated samples, while the AM-301-treated inserts had limited viral titer growth. In particular, at the end of the treatment period (Day 4), inserts that had received AM-301 had significantly lower viral titer (12- or 8-fold lower, respectively) than saline- or matrix-treated inserts. In the AM-301 no wash condition, inserts had a 12- or 15-fold lower titer than saline- or matrix-treated inserts, respectively. Similar results were seen in Protocol 2 (test substances applied at 30 h; Figure 4B): Day 4 samples treated with AM-301 had about 4- or 3-fold lower titer than saline- or matrix-treated inserts, respectively. At the same time point, the AM-301 no wash inserts had a 21- or 15-fold lower titer than the saline- and matrix-treated inserts, respectively. Viral titer for the inserts not exposed to SARS-CoV-2 was <231, indicating that the test substances and culture medium were free of viral contamination. Altogether, despite noticeable intragroup variability, these results suggest that AM-301 can mitigate an established infection even when applied only many hours after exposure to the virus.

**Figure 4.**
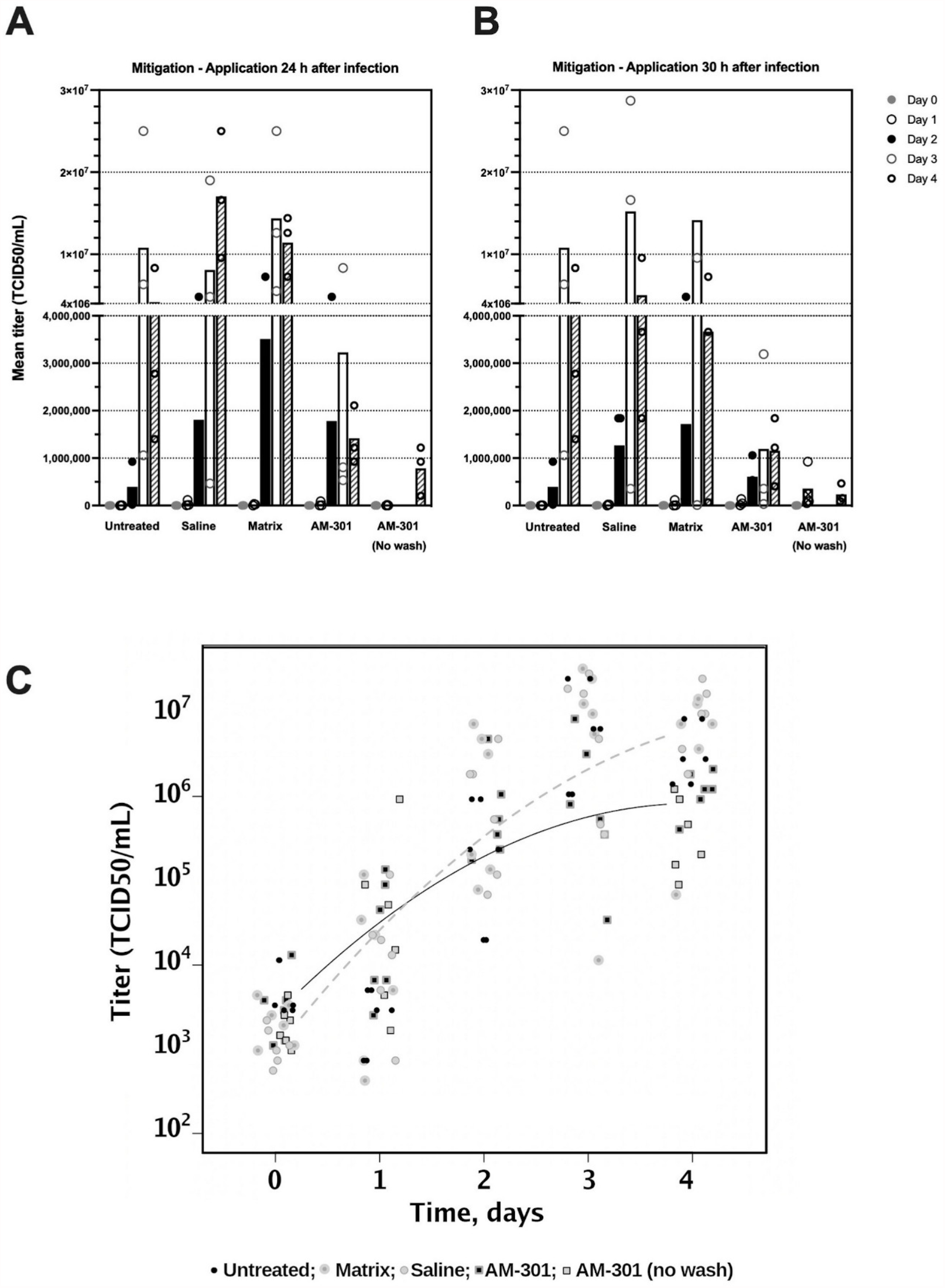
AM-301 in the mitigation of SARS-CoV-2 infection. **A, B** Bar charts with mean values and individual data points. Test substances were applied 24 or 30 h after the start of the experiment (Protocols 1 and 2, respectively). **C** Linear mixed-effect model. The log-linear scatter plot shows individual log-transformed data and concave curves for negative control samples (untreated, saline- and matrix-treated inserts; dashed curve) and for AM-301-treated inserts (continuous curve) (t=3.68, p<0.01). The model shows a deceleration in exponential growth, as is typically observed for sigmoidal behaviors.

Data from the mitigation experiment were analyzed with a linear mixed-effect model (Figure 4C), which revealed that: (i) the time profile of SARS-CoV-2 infection was not impacted by matrix or saline solution; (ii) the time profile of SARS-CoV-2 infection in the presence of AM-301, with or without the washing procedure (AM-301 and AM301 no wash, respectively), was significantly decelerated (t-value 3.68, p<0.01) compared to the untreated inserts and to inserts that received saline or matrix. Importantly, all experimental conditions were statistically equivalent at baseline (t-value 1.93), and there was no statistically significant difference between the application of AM-301 after 24 and 30 hours. All data were considered valid, and no outliers were excluded.

## Discussion

The nasal cavity, covered by pseudostratified ciliated epithelium and rich in mucus-secreting goblet cells, is the frontline barrier through which respiratory viruses, including SARS-CoV-2, pass to reach the cells of the lower airway [34,35]. The nose is not only a portal for infection, but also an important site of viral replication [15]. Therefore, the nose is an important target for prophylaxis and potentially also for therapy. SARS-CoV-2 cellular tropism in patients with early infection is restricted to the nasal ciliated cells of the oral squamous epithelium [11]. Nasal swabs from both symptomatic and asymptomatic patients contain higher viral loads than do throat swabs, with a consequentially higher possibility to transmit the virus through respiratory droplets [36–38].

To study the ability of AM-301 to prevent or reduce SARS-CoV-2 infection in the nasal mucosa, we used a well-established model of primary human nasal epithelium, MucilAir. AM-301 was studied to determine its compatibility with MucilAir, its efficacy in preventing MucilAir from being infected by SARS-CoV-2, and its ability to mitigate an established infection in MucilAir without any previous treatment. First, AM-301 and its matrix (lacking bentonite) had no detrimental effects on MucilAir inserts despite repeated application over 4 days: measures of tight junction integrity in treated cultures did not differ from those of untreated cultures, and an LDH assay of cytotoxicity revealed no increase in cell death. A slight reduction in cilia beating frequency was detected in AM-301 and matrix-treated inserts compared to the controls. However, there was no difference between AM-301 and its matrix, suggesting that bentonite per se is not responsible for this effect, which may rather be due to the viscosity of the preparation.

Pretreatment of MucilAir inserts with AM-301 (but not its matrix) was protective against SARS-CoV-2 infection, as just a daily application of the product led to a 2-log (99%) reduction in viral titer by Day 4. Inserts that received the product 24 or 30 h after viral infection also had a lower viral titer, corresponding to a 12- or 14-fold lower TCID50/mL at the end of the treatment. These data suggest that AM-301 may be effective as an aid to prophylaxis and to help reduce the viral titer in the upper respiratory tract as a treatment of a SARS-CoV-2 infection. The positive effects of AM-301 in reducing SARS-CoV-2 infection are supported by the observations from linear-mixed effect models, which showed significant decelerations in viral titer growth in both the prophylaxis and mitigation assays.

AM-301 was developed to mechanically prevent the virus from contacting the nasal mucosa and infecting the upper respiratory airways, and to trap or bind it for clearance by mucociliary clearance, ultimately helping to prevent a dramatic viral spread in the airways. Thus, AM-301 can be expected to lower the infectious viral load. This proposed mechanism is based on the hypothesis that charged viral particles interact with negative charges on cell membranes, making polyanionic substances a valid tool to impede this interaction [39,40]. The advantage of this type of non-pharmacological approach is the broad spectrum of action on viruses and allergens, which can be trapped as soon as they enter the body passing through the nose.

AM-301 (Bentrio) is a medical device containing only excipients and inert ingredients generally recognized as safe. Clays such as bentonite, a key component of the formulation, can also be used to detoxify water of fluoride or heavy metals [21], and bentonite is the basis of an oral treatment for acute infectious diarrhea (diosmectite, Smecta, available in some countries [41]). Its virus-capturing properties have been known for many years, but to our knowledge this is its first application in capturing airborne viruses.

Bentonite confers thixotropic properties to AM-301, permitting its easy application with a nasal spray pump, which results in a protective film once it contacts the nasal epithelium. The lack of preservatives, decongestants and other pharmacologically active molecules was intended to ensure maximal safety and compatibility with the nasal microbiome. This feature is particularly important, since a reduction in nasal microbiota diversity seems to be associated with pathological conditions of the respiratory tract, such as asthma, bronchiolitis and flu, and has an influence on the immune response in chronic rhinosinusitis [42]. Therefore, the use of AM-301 for long periods of time should not alter the natural properties of the nasal epithelial barrier [43], but rather should endow it with an additional feature crucial for the maintenance of tissue homeostasis and for the defense from pathogens, toxins, allergens and pollutants.

Conducting this study was challenged by the several factors that may account for the noticeable intragroup variability: differences in the mucus quantities produced by the MucilAir inserts, and technical difficulties in the washing procedures to evenly remove the mucus from all inserts. However, these in vitro results highlight the potential and relevance of AM-301, since the MucilAir tissue model can be considered a worst-case scenario of the in vivo situation. MucilAir lacks an immune system to protect against infection, and mucociliary clearance of viral particles does not occur. AM-301 was applied once every 24 h instead of 2-3 times per day as it might be used by people. The encouraging results of this study call for further investigations in vivo and in humans, to further evaluate AM-301 as a medical device with a broad spectrum of action against a battery of viruses, allergens and pollutants.

Intriguingly, since nasal spray components not absorbed by the nasal mucosa are discharged from the nose to the pharynx, AM-301 may exert its beneficial virus-capturing effects also in the throat. Since the oral cavity is where the highest production of aerosols and droplets occurs [44,45], this action might decrease the potential for viral transmission to other people by exhalation, talking or coughing [46]. The ability of AM-301 to reduce viral transmission is currently under investigation.

Nowadays, the entire world must deal with the health, social and economic damage that the COVID-19 pandemic caused. Several therapies and vaccines have been developed to allow us to go back to normal life. The ability of SARS-CoV-2 to mutate quickly and the high cost of medications, vaccines and protective measures as well as supply chain challenges make this process more difficult in developing countries. Viral epidemics and pandemics are expected to become more frequent due to population increases, urbanization, warmer climates and, especially, increases in the frequency and diversity of wildlife–livestock–human interfaces [47]. It is thus critical to have efficacious strategies that can contrast viral spread rapidly and that are sufficiently unspecific to be able to contrast different viral strains.

AM-301 is a safe, non-pharmacological, easy-to-use nasal spray that can reduce the risk of infection from SARS-CoV-2 and potentially from other viruses by acting as an “intranasal mask”. It appears to be well suited for self-protection and as a complement to other preventive measures such as increased hygiene, social distancing, and vaccination. Further, its stability at high temperature, tested and confirmed in accordance with relevant guidelines, allows its use in warm climates and developing countries where protection from airborne viruses is challenging due to costs or availability of more sophisticated forms of protective devices.

## Author contributions

FF and RJ conceived and managed the experiments, SM carried out the efficacy experiments, VF contributed by interpreting the results, writing, and editing the manuscript, MB did the statistical modelling.

## Acknowledgments

Valerie Matarese provided writing and editing services on early versions of this manuscript.

